# Ferroptosis Inhibition Combats Metabolic Derangements and Improves Cardiac Function in Pulmonary Artery Banded Pigs

**DOI:** 10.1101/2024.04.24.590907

**Authors:** Felipe Kazmirczak, Ryan Moon, Neal T. Vogel, Walt Tollison, Matt T.Lahti, John P. Carney, Jenna B Mendelson, Todd Markowski, LeeAnn Higgins, Kevin Murray, Candace Guerrero, Kurt W. Prins

**Author notes:** Corresponding Author Kurt W. Prins, MD, PhD, Associate Professor of Medicine, Director of Translational Research in Pulmonary Hypertension and Right Heart Failure, 2231 6^th^ St SE, Minneapolis, MN.

## Abstract

Right heart failure (RHF) is a leading cause of mortality in multiple cardiovascular diseases and preclinical and human data suggest impaired metabolism is a significant contributor to right-sided cardiac dysfunction. Ferroptosis is a nonapopotic form of cell death driven by impaired metabolism. Rodent data suggests ferroptosis inhibition can restore mitochondrial electron transport chain function and enhance cardiac contractility in left heart failure models, but the effects of ferroptosis inhibition in translational large animal models of RHF are unknown. Here, we showed ferrostatin-1 mediated ferroptosis antagonism improve right heart structure and function in pulmonary artery banded pigs. Molecularly, ferrostatin-1 restored mitochondrial cristae structure and combatted downregulation of electron transport chain proteins. Metabolomics and lipidomics analyses revealed ferrostatin-1 improved fatty acid metabolism. Thus, these translational data suggest ferroptosis may be a therapeutic target for RHF.

**Graphical Abstract:** 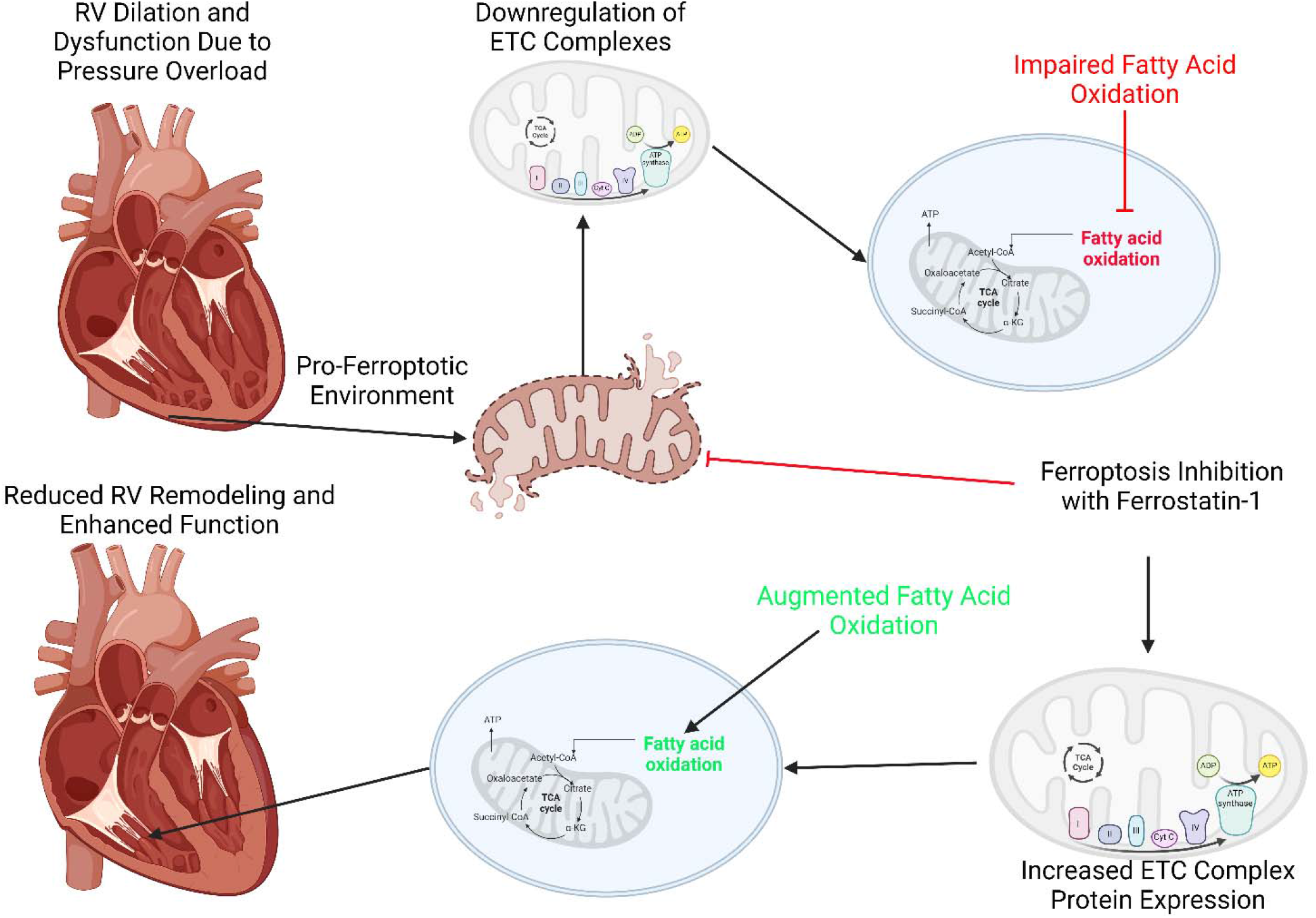

## To the Editor

Right ventricular failure (RVF) is the leading cause of morbidity and mortality in pulmonary arterial hypertension (PAH), but we currently lack therapies that enhance right ventricular function(1). Both rodent and human data suggest mitochondrial dysfunction, marked by impairments in fatty acid oxidation and alterations in electron transport chain homeostasis, drives RVF(1–3). While fatty acid oxidation is the leading source of ATP generation in the heart, disruptions in fatty acid oxidation can promote ferroptosis, a lipid peroxidation-dependent mode of cell death(4). Ferroptosis also suppresses electron transport chain function and compromises cardiac function in left ventricular failure(5). Rodent and human data suggest ferroptosis plays a role in PAH-mediated RVF. In rodent RVF induced by monocrotaline, intermittent fasting suppresses the upregulation of multiple pro-ferroptotic proteins, which is correlated with augmented RV contractility(6).

Transcriptomics data show heme oxygenase 1, a pro-ferroptotic protein, is upregulated in rodents and humans with PAH-associated RVF(2, 7). Moreover, ferrostatin-1, a small molecule ferroptosis inhibitor, enhances RV function in monocrotaline rats(8, 9), but that is in the setting of reduced afterload. Thus, the direct physiological and molecular effects of ferroptosis inhibition on the RV are less well described. Moreover, the potential benefits of ferroptosis suppression in a translational large animal model of RVF are unknown.

Here, we addressed these knowledge gaps by evaluating the effects of ferroptosis inhibition in a porcine model of RVF induced by pulmonary artery banding (PAB)(10). To improve translatability, ferrostatin-1 (0.2 mg/kg diluted in 2% dimethyl sulfoxide, 50% polyethylene glycol, 5% Tween 80, and 43% double distilled water), treatment via intramuscular injection began three weeks after PAB to allow for the development of right ventricular dysfunction. PAB pigs treated with an equivalent volume of ferrostatin-1 vehicle (PAB-Placebo) were also evaluated. End-point analyses were performed six weeks after PAB. Primary end-point was right ventricular ejection fraction (RVEF) as determined by cardiac magnetic resonance imaging (cMRI), and blinded analysis was performed by FK. Right heart catheterization probed RV pressures/afterload, quantitative proteomics of mitochondrial enrichments evaluated mitochondrial protein regulation(6), and a combined metabolomics/lipidomics analysis (Biocrates MxPQuant 500 XL kit) quantified >1200 metabolites/lipids(9) in RV free wall specimens. Data quality analysis/outlier detection of proteomics/metabolomics data included inspection of hierarchical cluster analysis. Animal studies were approved by the University of Minnesota Institutional Animal Care and Use Committee (2112-39666A).

Ferrostatin-1 exerted significant therapeutic effects as cMRI revealed ferrostatin-1 increased RVEF by 11% (**Figure 1 A and B**). In addition, ferrostatin-1 reduced pathological RV dilation (**Figure 1C**). Importantly, these changes were not due to alterations in RV workload as the anatomical (% pulmonary artery stenosis, PAB-Placebo: 88±0.8% PAB-Fer-1: 87±1.9%, *p*=0.61, unpaired t-test) and RV systolic pressure (Control: 21±2 PAB-Placebo: 57±3 PAB-Fer-1: 50±5, *p*=0.19 comparing PAB-Placebo versus PAB-Fer-1 using one-way *ANOVA* with Tukey’s multiple comparison test) were not significantly different between PAB-Placebo and PAB-Fer-1. Ferrostatin-1-mediated corrections in RV remodeling and function counteracted PAB-induced right atrial dilation (Right atrial end-diastolic volume indexed to body mass: Control: 1.3±0.1, PAB-Placebo: 2.1±0.2, PAB-Fer-1: 1.3±0.1, *p*=0.002 PAB-Placebo versus PAB-Fer-1, *p*-values determined by one-way ANOVA with Tukey’s multiple comparisons test), and right atrial ejection fraction (**Figure 1D**). Finally, ferrostatin-1 nonsignificantly reduced late gadolinium enhancement at the RV insertion sites (Table 1 and Supplemental Figure 1).

**Table 1:**
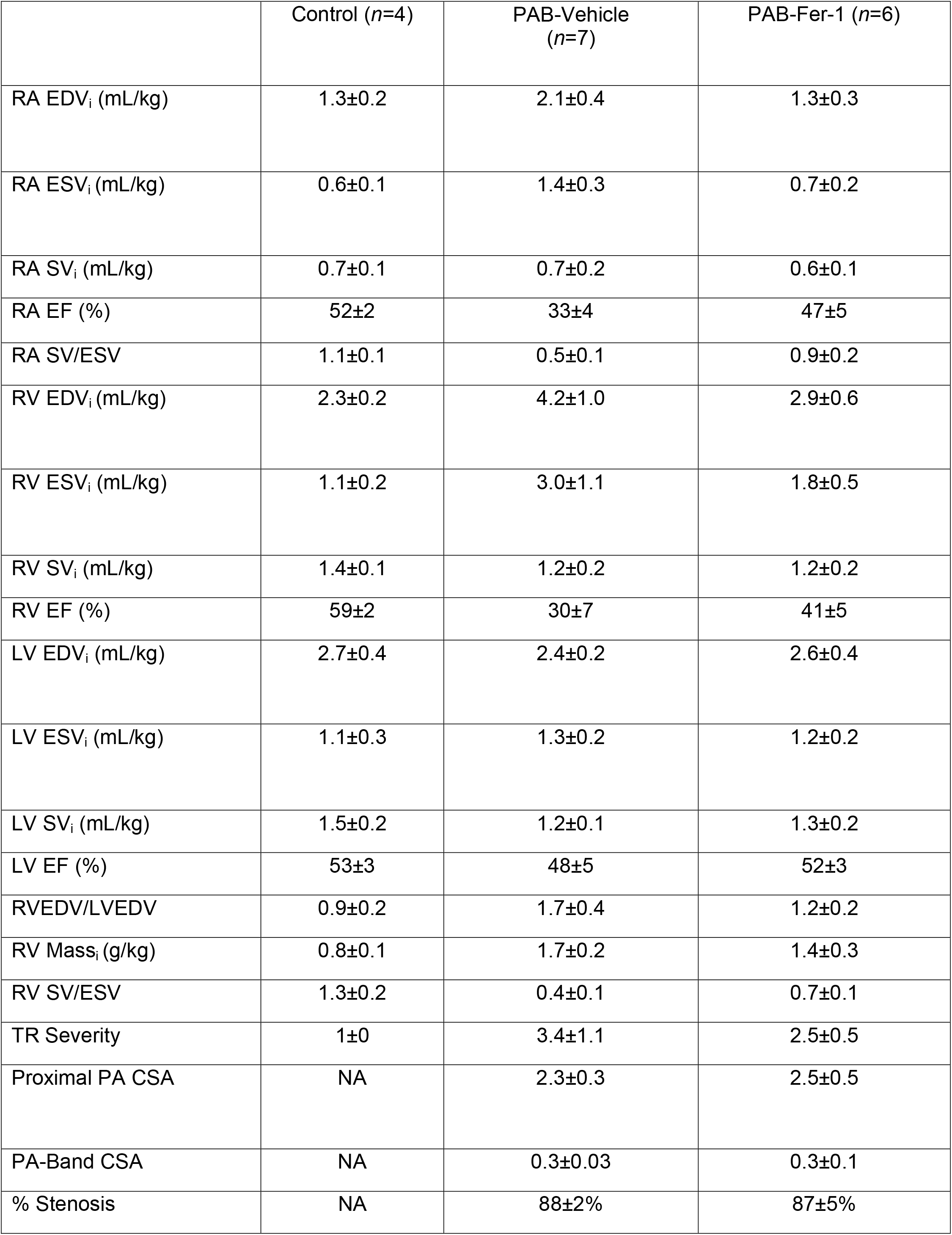
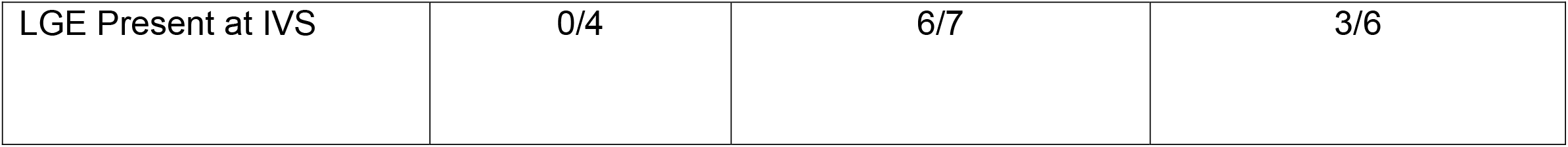
Comprehensive Cardiac MRI Evaluation.

**Figure 1:**
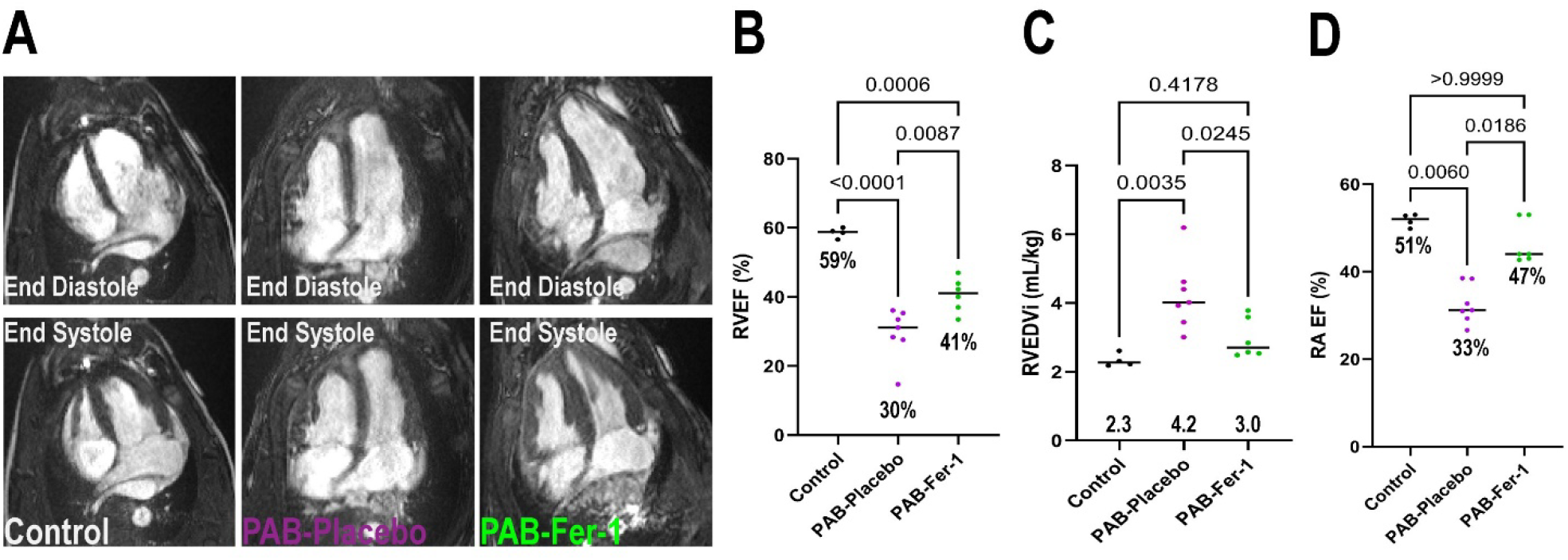
Ferrostatin-1 Treatment Improves Right Ventricular Structure and Function in Pulmonary Artery Banded Pigs. (A) Representative four-chamber cardiac magnetic resonance images in end-systole and end-diastole of control PAB-Vehicle, and PAB-Fer-1 animals. (B) Blinded quantification of right ventricular ejection fractions, *p*-values determined by one-way *ANOVA* with Tukey’s multiple comparison test. Ferroptosis inhibition counteracted RV dilation (C), *p*-values determined by one-way *ANOVA* with Tukey’s multiple comparison test and increased right atrial ejection fraction (D), *p*-values determined by Kruskal-Wallis test and Dunn’s multiple comparisons test.

Because ferroptosis modulates mitochondrial homeostasis and electron transport chain function(5), we evaluated RV mitochondrial morphology and protein regulation. Electron microscopy revealed ferrostatin-1 restored cristae integrity (Supplemental Figure 2). RV mitochondrial proteomics analyses suggested ferroptosis inhibition altered the overall molecular signature as PAB-Fer-1 animals clustered closer to control than PAB-Placebo as demonstrated by hierarchical cluster analysis (**Figure 2A**). KEGG pathway analysis of the 250 proteins most important for differentiating the three experimental groups as determined by partial least squares discriminant analysis identified oxidative phosphorylation as the most enriched pathway (**Figure 2B**). Nearly all oxidative phosphorylation proteins were reduced in the PAB-Placebo group, but ferrostatin-1 treatment partially rescued the downregulation of many oxidative phosphorylation proteins (**Figure 2B**). Then, we probed the RV metabolic consequences of impaired mitochondrial protein regulation with metabolomics/lipidomics analysis. The global RV metabolomics/lipidomic signature was changed when PAB-Placebo animals were compared to control (**Figure 2C**). However, PAB-Fer-1 animals appeared to be an intermediate between control and PAB-Placebo (**Figure 2C**). Random forest classification identified the 15 metabolites/lipids that were divergent comparing the three groups, and changes in phosphatidylethanolamines and metabolites of fatty acid metabolism [diacylglycerides and acylcarnitines] were among the most important variables that differentiated the three groups. Therefore, we evaluated fatty acid metabolites and we found ferrostatin-1 partially mitigated the reduction in triacylglycerides, diacylglycerides, and acylcarnitines (**Figure 2C**).

**Figure 2:**
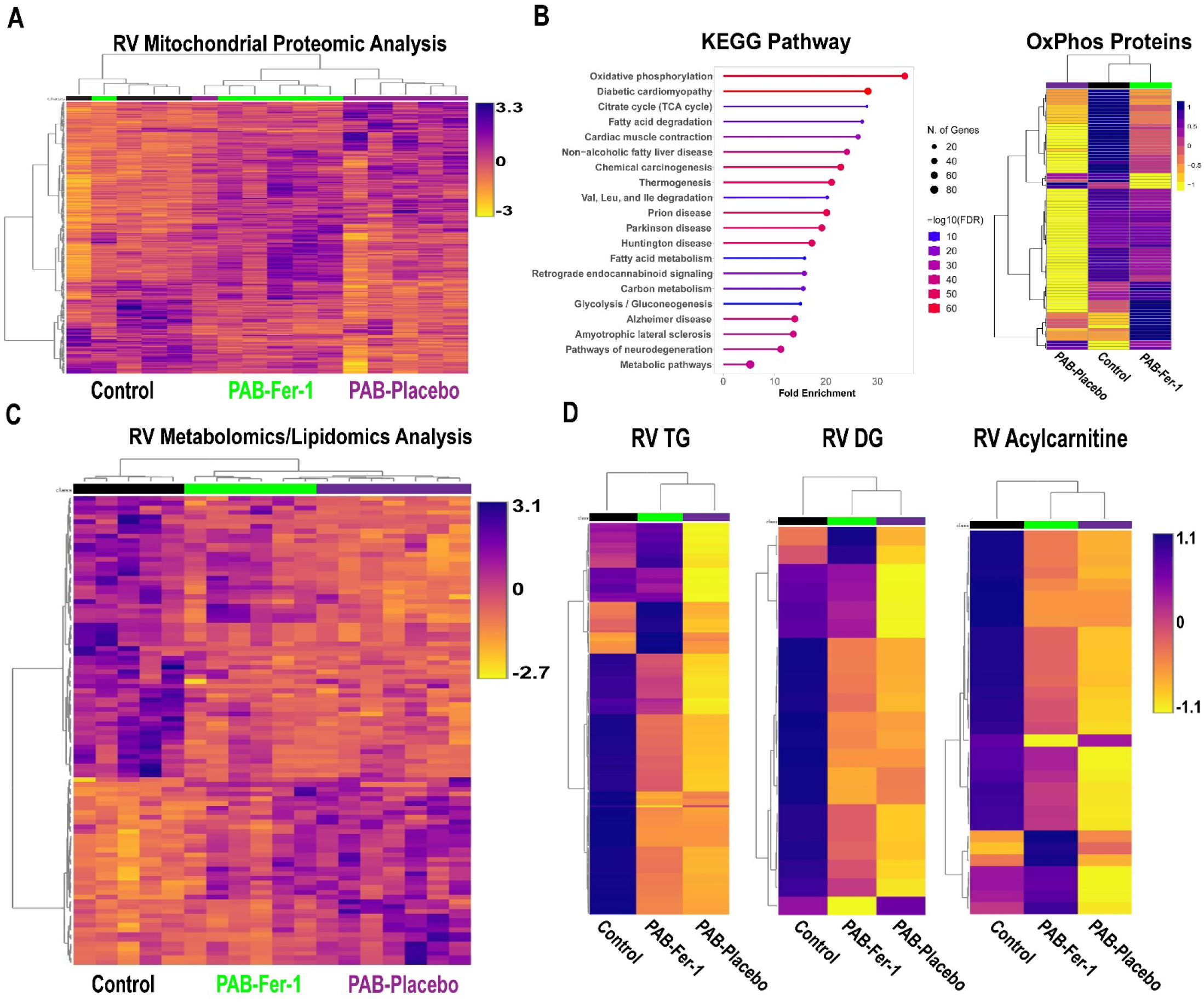
Ferrostatin-1 Counteracts Mitochondrial Metabolic Dysregulation in the RV of PAB Pigs. (A) Hierarchical cluster analysis of mitochondrial enrichments. (B) KEGG pathway analysis of the top 250 proteins that differentiated the three groups (left) and hierarchical cluster analysis of protein subunit abundances of oxidative phosphorylation proteins (right). (C) Hierarchical cluster analysis of total metabolites/lipids in RV specimens. (D) Hierarchical cluster analysis of triacylglycerides (TG), diacylglycerides (DG), and acylcarnitines. Ferroptosis inhibition partially prevented the reduction of TG, DG, and acylcarntines in RV samples.

Our study has important limitations that we acknowledge. First, it was conducted only in castrated males and thus biological sex was not evaluated. Second, our lipidomic analysis did not detect lipid peroxidation events, which are technically challenging because lipid peroxides are quite labile and even when samples are stored at −80 °C signal is lost(11). Because these experiments were conducted over months, we believed a lipid peroxidation assessment was not technically feasible. We observed more significant downregulation of mitochondrial metabolic proteins in this PAB model than our previous study(10), however these pigs had more compromised RV function (RVEF:30% versus 38%). A 5% change in RVEF has clinically meaningful implications(12), and thus there are likely alterations in the molecular signature as RVEF drops, which may account for some of the differences. Our data demonstrated triacylglycerides were reduced in PAB-Placebo RVs, which is the opposite of what PAH patients exhibit(13). There are several potential explanations for these discrepancies including inherent species differences in triacylglyceride metabolism, caloric intake and dietary composition, which modulates RV triacylglyceride storage(14), and methodological and site of evaluation differences. We evaluated triaglycerides in RV free wall specimens using lipidomics while Brittain *et al*., quantified triacylglycerides with proton spectroscopy in the intraventricular septum(13). Undoubtedly, further studies are needed to clarify and understand these opposing results. We excluded two PAB-Placebo pigs because of blunted growth, which could both reduce afterload severity over the time course of the experiment and indicate an undefined systemic process. Finally, one PAB-Fer-1 animal died from an incarcerated hernia, and autopsy revealed peritonitis, perforated intestine, but no pleural effusion, suggesting RV failure was not a significant contributor.

In conclusion, our study demonstrates ferroptosis inhibition combats mitochondrial metabolic protein downregulation and disrupted fatty acid metabolism. These molecular changes are paired with significant improvements in right heart structure and function. Because we recently demonstrated ferroptosis inhibition counteracts pulmonary vascular remodeling in rodents(9); we propose ferroptosis inhibition may have advantageous effects for the entire RV-PA unit, which makes it an attractive therapeutic target for PAH.

## Methods

### Animal Groups and Treatment

8-10 week old Yorkshire Cross castrated male pigs were randomized into two treatment groups: ferrostatin-1 (PAB-Fer-1) or Placebo (PAB-Placebo). To improve translatability, vehicle (2% dimethyl sulfoxide, 50% polyethylene glycol, 5% Tween 80, and 43% double distilled water) or ferrostatin-1 (0.2 mg/kg, daily intramuscular injection) treatment started three weeks after PAB and continued for a total of three weeks. Five control pigs were purchased from vendor and housed at the University of Minnesota without any interventions. Animals studies were approved by the University of Minnesota Institutional Animal Care and Use Committee protocol number 2112-39666A. One control animal suffered a pericardial effusion during hemodynamic study so no cardiac MRI was performed. Two PAB-Vehicle animals were excluded: both due to poor growth with having poor weight gain during the experimental timeline and one having insufficient PAB severity. One PAB-Fer-1 animal died before end-point analysis and autopsy revealed it suffered a hernia, which was deemed to be the cause of death.

### Pulmonary Artery Banding Procedure

Animals were initially sedated with Telazol (4 mg/kg) and Xylazine (0.8mg/kg) via intramuscular injection. After allowing the drugs to take effect, an ear catheter was placed for intravenous access. The IV catheter was then connected to an IV infusion of 0.9% Normal saline (NaCl).

Anesthesia was induced by administering 1-2 mg/kg propofol IV to effect. Following evaluation for an appropriate depth of anesthesia, the animals were intubated with an appropriately sized endotracheal tube. Ceftiofur (5 mg/kg, IM) was administered once during induction for prophylactic antibiotic coverage.

When confirmed the animal reached a deep plane of anesthesia, 1.1 mg/kg succinylcholine chloride was administered IV. A left thoracotomy was performed through the 4^th^ intercostal space. The pericardium was opened parallel and anterior to the phrenic nerve and cradled with 2-0 braided polyester free ties. The main pulmonary artery was isolated proximal to the bifurcation. An umbilical tape was double looped around the pulmonary artery using a right angle. The umbilical tape was tightened, creating an hour glass shape, while the vital signs were monitored. When the animal appeared to be stable, a 19 gauge needle on a fluid filled pressure monitoring line was placed in the RV. The post-band RV pressure was recorded. Allowing up to 15 minutes ensuring the animal remained stable, the umbilical tape was tied and a clip was placed to secure the knot. The pericardium was loosely approximated, and the chest cavity was washed with a warm antibiotic solution. A chest tube was then placed in the chest through an intercostal space caudal to the thoracotomy, exteriorized through the skin and connected to a water sealed vacuum drainage reservoir. The rib, muscle, and skin layers were closed in standard fashion.

The animas were observed during the immediate post-operative period for bleeding. The animal were weaned from the ventilator to initiate spontaneous breathing. Supplemental oxygen was provided through the endotracheal tube. When the animal was breathing normally, substantial fluid was no longer draining from the chest tube, and negative thoracic pressure was established, the chest tube was removed. Once the animal was determined to be stable, it was moved to RAR post-operative care area. The endotracheal tube was removed when appropriate. Post-operative antibiotic (Clavamox 14 mg/kg BID for seven days) following the main pulmonary artery banding procedure was administered.

### Hemodynamic study

The animals were initially sedated with Telazol (4 mg/kg) and Xylazine (0.8mg/kg) via IM delivery. After allowing the drugs to take effect, an ear catheter was placed for intravenous access. The IV catheter was then connected to an IV drip with 0.9% Normal saline (NaCl). Anesthesia was induced by administering 1-2 mg/kg propofol IV. Following evaluation for an appropriate depth of anesthesia, the animals were intubated with an appropriately sized endotracheal tube. The animals were placed on a table in the right lateral decubitus position. Mechanical ventilation was initiated at 10-15 breaths per minute at a tidal volume of 1.5 x body weight (kg), oxygen at 3 L/min, and isoflurane set between 1 and 3%, as needed, to maintain anesthesia. Vital signs were monitored (ECG, SpO2, EtCO2, temperature). When confirmed the animal reached a deep plane of anesthesia, arterial and venous interventional access was gained through the jugular vein and femoral artery. The artery and vein were accessed via a surgical cut down. Heparin (250 mg/kg, IV) was administered prior to introducing a vascular sheath. An appropriately sized introducer sheath was placed in the jugular vein and femoral artery lumen and secured. A pressure line was connected to the femoral artery introducer for systemic pressure monitoring. A 7F Swan-Ganz catheter was placed in the jugular vein\ introducer port to obtain hemodynamic data. The systemic artery pressure and right sided pressures were connected pressure transducers, which were displayed on AD instruments LabChart.

### Cardiac magnetic resonance imaging/angiography (MRI/MRA) examination

All studies were performed after animals had hemodynamic study and thus all animals were intubated and sedated as described above. Studies were performed at the University of Minnesota using a Siemens 1.5 Tesla AERA scanner (Siemens, Malvern, PA) with phased-array coil systems. The examination included localizers to assess cardiac position and a standard segmented steady-state free-precession cine sequence to assess cardiac volumes and function. The imaging parameters were as followed: typical repetition time of 3.0–3.5 milliseconds, echo time of 1.2–1.5 milliseconds, in-plane spatial resolution of 1.8×1.4 millimeters, and temporal resolution of 35–40 milliseconds. Short-axis images were acquired with a slice thickness of 6 millimeters from the roof of the right atrium to the apex of the left ventricle. Long axis cines were obtained in the four-chamber, three-chamber and two-chamber views with dedicated two-chamber RV view. The cardiac magnetic resonance examination sequences were gated with electrocardiogram. Following this, contrast-enhanced magnetic resonance angiography was performed with contrast bolus timed to trigger at main pulmonary artery to evaluate the pulmonary artery banding severity. CMR analyses were performed using standard software (Precession by Heart Imaging Technologies, Durham, NC). Left and right atrial and ventricular end-diastolic and end-systolic volumes, ejection fractions, and mass were quantified by planimetry of the end-diastolic and end-systolic endocardial and end-diastolic epicardial borders on the short-axis cine images. All CMRI analyses were performed blindly by FK.

### Confocal Microscopy

Confocal microscopy was performed on RV free wall sections from all animals included in the final analysis to evaluate RV cardiomyocyte hypertrophy. Briefly, RV free wall slides were de-paraffinized through xylene and ethanol washes, heated in Reveal Decloaker Buffer (BioCare Medical, Pacheco, CA) for 30 minutes. Sections were washed/blocked with 5% goat serum in PBS and then and stained WGA-488 at a dilution of 1:50 for 30 minutes at 37°C. Samples were washed with PBS, exposed to Hoechst stain, and treated with an autofluorescence quenching kit (Vector Laboratories). Finally, samples were mounted in Prolong Glass Antifade Mountant (Thermo Scientific). Images were collected on a Zeiss LSM 900 Airyscan 2.0 microscope and blindly analyzed by RM. RV cardiomyocyte diameter was determined using FIJI (NIH).

### Mitochondrial Ultrastructural Analysis

RV free wall tissue was placed into fixative (4% paraformaldehyde + 1% glutaraldehyde in 0.1M phosphate buffered, pH 7.2 (PB)). After fixation, tissue was washed with PB, stained with 1% osmium tetroxide, washed in H_2_O, stained in 2% uranyl acetate, washed in H2O, dehydrated through a graded series of ethanol and acetone and embedded in Embed 812 resin. Following a 24 hour polymerization at 60°C, 0.1 µM ultrathin sections were prepared and post-stained with lead citrate. Micrographs were acquired using a JEOL 1400 Plus transmission electron microscope (JEOL, Inc., Peabody, MA) at 80 kV equipped with a Gatan Orius camera (Gatan, Inc., Warrendale, PA) at the Mayo Clinic. RV mitochondrial morphology was analyzed by RM, and cristae morphology scoring was performed as described(1).

### Quantitative Mitochondrial Proteomics

TMT-16plex quantitative proteomics, performed at the University of Minnesota Center for Metabolomics and Proteomics, of RV mitochondrial enrichments (Abcam) evaluated RV mitochondrial protein regulation in *n*=4 control, *n*=6 PAB-Vehicle, and *n*=6 PAB-Fer-1 specimens as we have previously described (2–4). Protein abundance was quantified using Proteome Discoverer Software version 3.0 and used for statistical analysis as described below.

### RV Metabolomics/Lipidomics Analysis

Global metabolomics analysis of frozen right ventricular free wall specimens on *n*=5 Control, *n*=7 PAB-Placebo, and *n*=6 PAB-Ferrostatin-1 was performed at the University of Minnesota Center for Metabolomics and Proteomics using the Biocrates’ MxP® Quant 500 kit. Approximately 100 mg of RV free wall was placed in 2.0 mL Precelly standard tubes and homogenized 3 times for 30 seconds at 5,800 rpm. Samples were centrifuged at 10,000*g* for 5 minutes at 4°C and supernatant was collected. 10µL of the extract was loaded onto a well insert. Classes of metabolites were determined using 50µL of 1:1:1:0.16 water:EtOH:pyridine:phenyl isothiocyanate solution and incubated for an hour. ABSciex QTRAP 5500 triple-quadrupole (Farmington, MA, USA) mass spectrometer was used to perform metabolomic assays.

### Statistical Analysis

The primary end-point of this study was right ventricular ejection fraction (RVEF). We estimated a sample size of *n*=10 animals to detect a significant difference in RVEF, however interim analysis revealed the primary end-point was met so the study was terminated. Statistical analyses were performed using Prism 10.0 (GraphPad Software). Normality of data was determined using the Shapiro-Wilk test. If data were normally distributed and there was equal variance as determined using the Brown-Forsythe test, One-way analysis of variance with Tukey’s multiple-comparisons test was performed. If there was unequal variance, Brown-Forsythe and Welch analysis of variance with Dunnett multiple-comparisons test was completed. If the data were not normally distributed, the Kruskal-Wallis test and Dunn’s multiple-comparisons test were used when comparing three groups. *P*-values of <0.05 were considered to indicate statistical significance. Hierarchical cluster analyses, partial least squares discriminate analysis, and random forest classification of proteomics and metabolomics/lipidomics data were performed using MetaboAnalyst software. The relative abundance of each protein in the proteomics experiments were determined using Proteome Discover Software as we have previously described(2). Kyoto Encyclopedia of Genes and Genomes (KEGG) pathway analysis of the top 250 proteins most important for distinguishing the three experiment groups as determined by Random Forest Classification was performed using ShinyGO 0.76.3 (http://bioinformatics.sdstate.edu/go/). All pathways enriched were defined by a false discovery rate of *p*<0.05.

## Data Availability

All data will be provided upon reasonable request. The raw proteomics (10.6084/m9.figshare.24878304) and metabolomics/lipidomics (10.6084/m9.figshare.24943449) data are currently available on Figshare.com with the listed DOI.

---

**Supplemental Figure 1:**
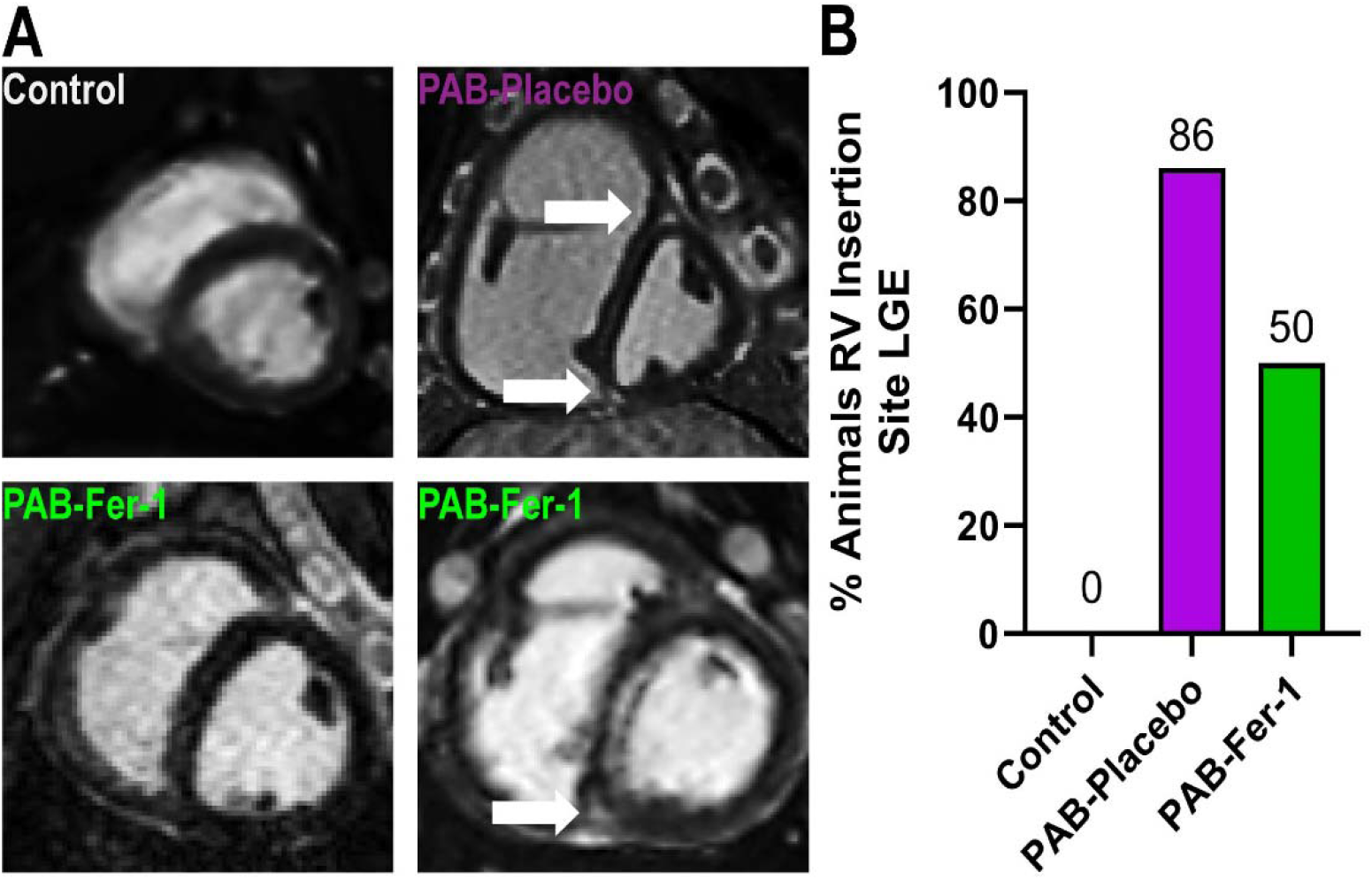
Ferrostatin-1 partially mitigated RV insertion site late gadolinium enhancement. (A) Representative cMRI images of LGE. Arrows indicate RV insertion site LGE. (B) Proportion of animals with RV LGE.

**Supplemental Figure 2:**
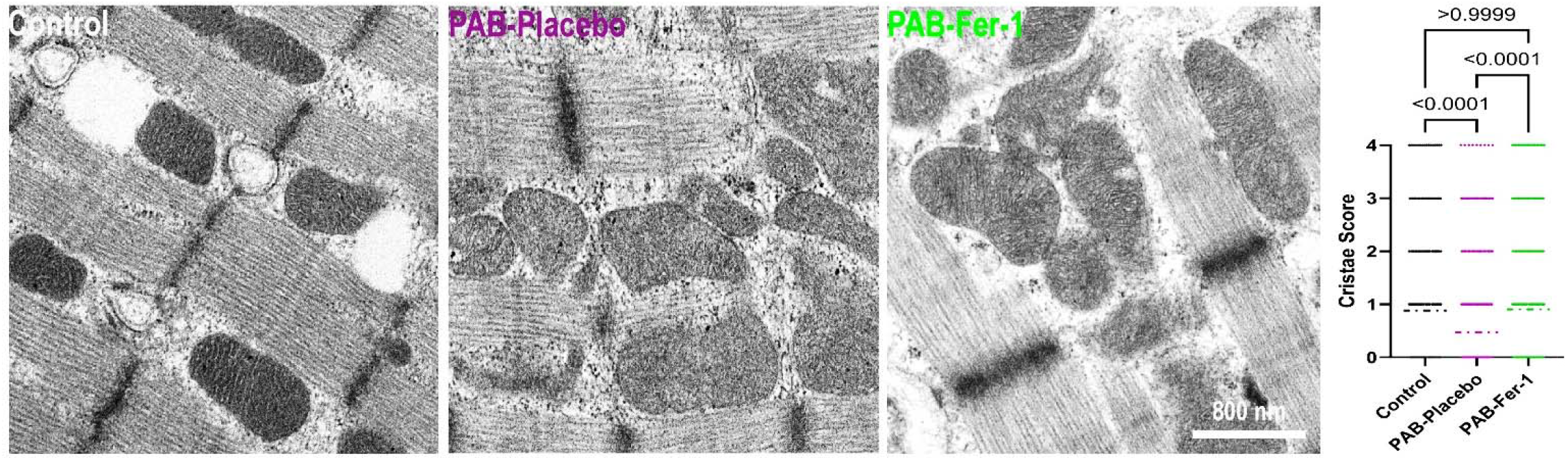
Ferrostatin-1 Improved Mitochondrial Cristae Morphology. Representative electron micrographs of RV cardiomyocytes depicting mitochondrial cristae structure. Blinded quantification of mitochondrial cristae morphology. Ferrostatin-1 improved cristae morphology.

**Supplemental Figure 3:**
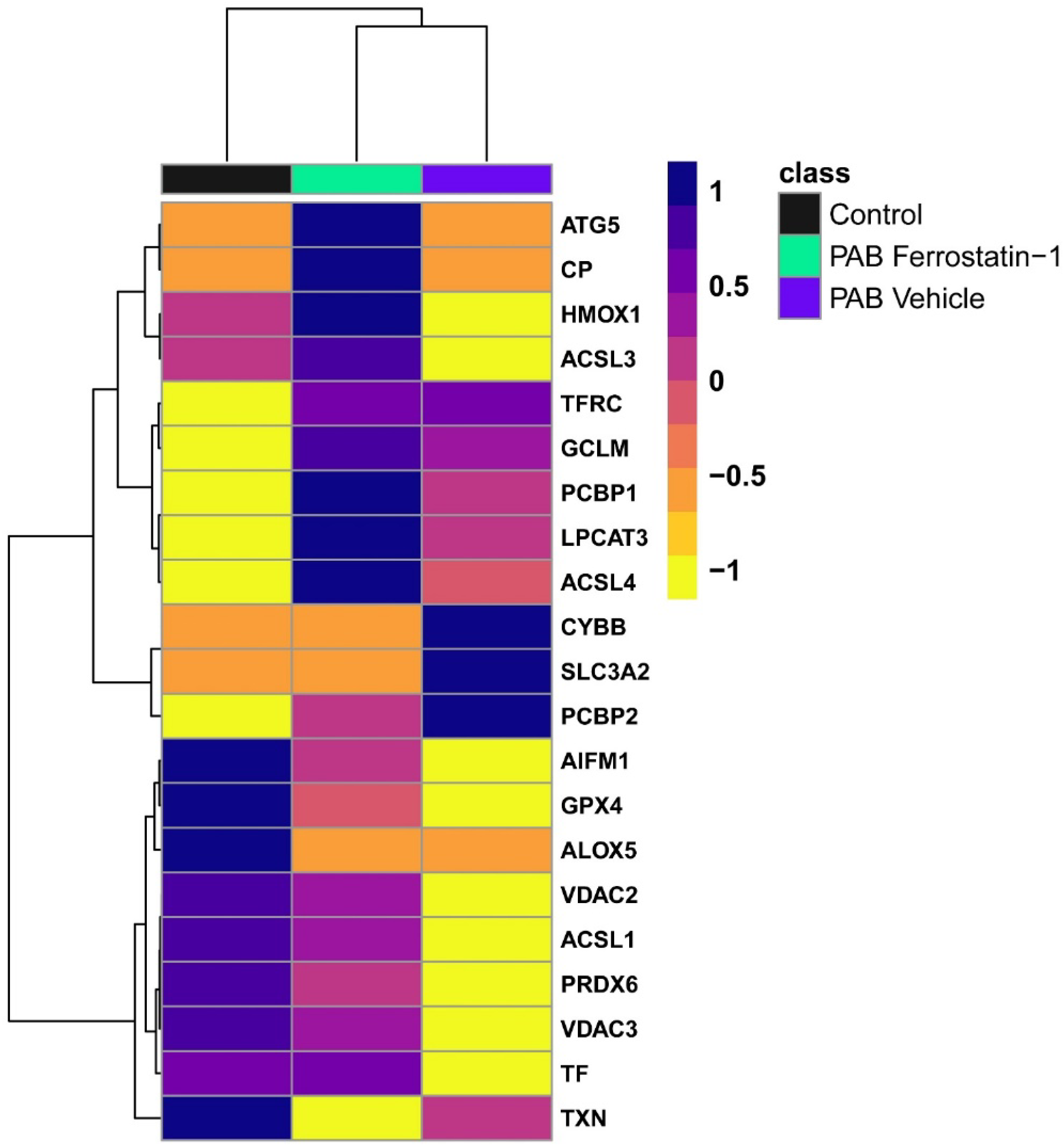
Hierarchical cluster analysis of ferroptosis proteins.

**Supplemental Figure 4.**
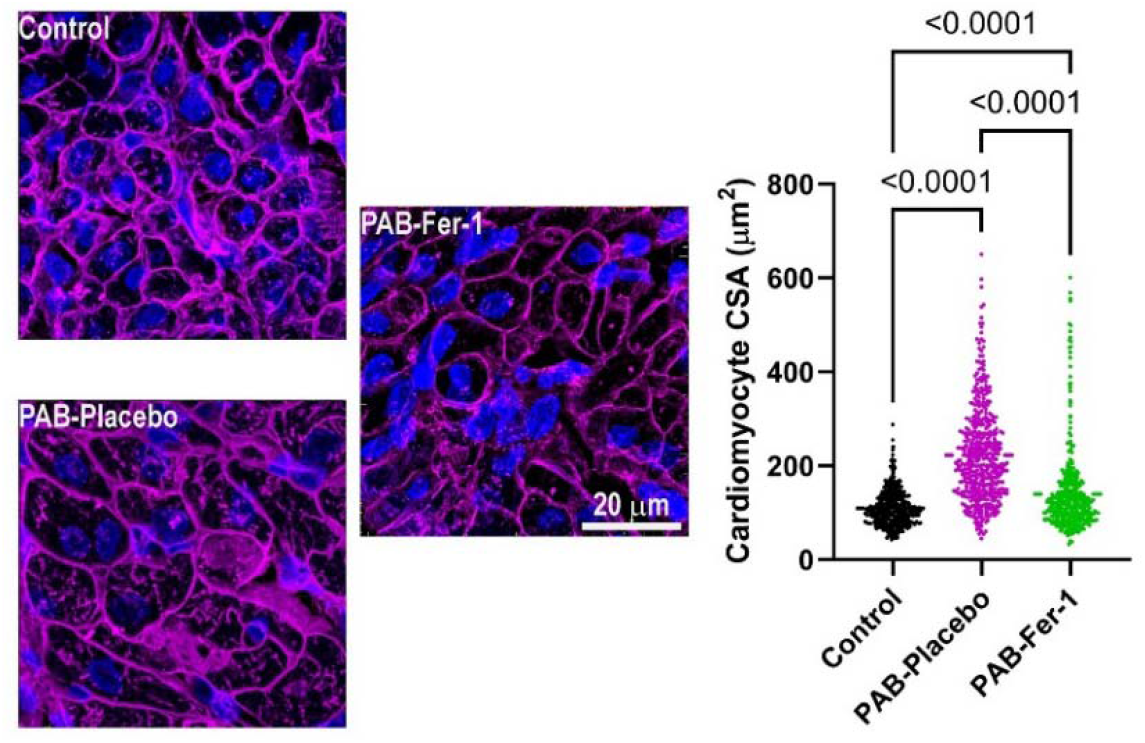
Representative confocal micrographs stained with wheat germ agglutinin (purple) and DAPI (blue) staining of RV cross sections and subsequent quantification of cardiomyocyte cross-section area

