## Supplemental Data for "Ferroptosis Inhibition Combats Metabolic Derangements and Improves Cardiac Function in Pulmonary Artery Banded Pigs"

Supplemental Figure 1: Ferrostatin-1 counteracts pathological changes in the right heart despite no changes in RV afterload. Cardiac MRI examination revealed ferrostatin-1 reduced right ventricular end-diastolic volume indexed to body mass (A), the ratio of right ventricular end-diastolic volume to left ventricular end-diastolic volume (B), and tricuspid regurgitation severity (C). (D) Representative confocal micrographs stained with wheat germ agglutinin (purple) and DAPI (blue) staining of RV cross sections and subsequent quantification of cardiomyocyte cross-section area (E). (F) Three-dimensional reconstruction (above) and two-dimensional views of pulmonary artery magnetic resonance angiogram at the level of the aortic valve. (G) Quantification of the percent of pulmonary artery stenosis in PAB-Placebo and PAB-Fer-1 animals. (H) Hemodynamic evaluation of RV afterload as defined by right ventricular systolic pressure relative to systemic arterial systolic pressure. Ferroptosis inhibition mitigated pathological right atrial remodeling and dysfunction as right atrial end-diastolic volume relative to body weight (I), right atrial ejection fraction (J), and estimated right atrial-right ventricular coupling (K) were all improved with ferrostatin-1.

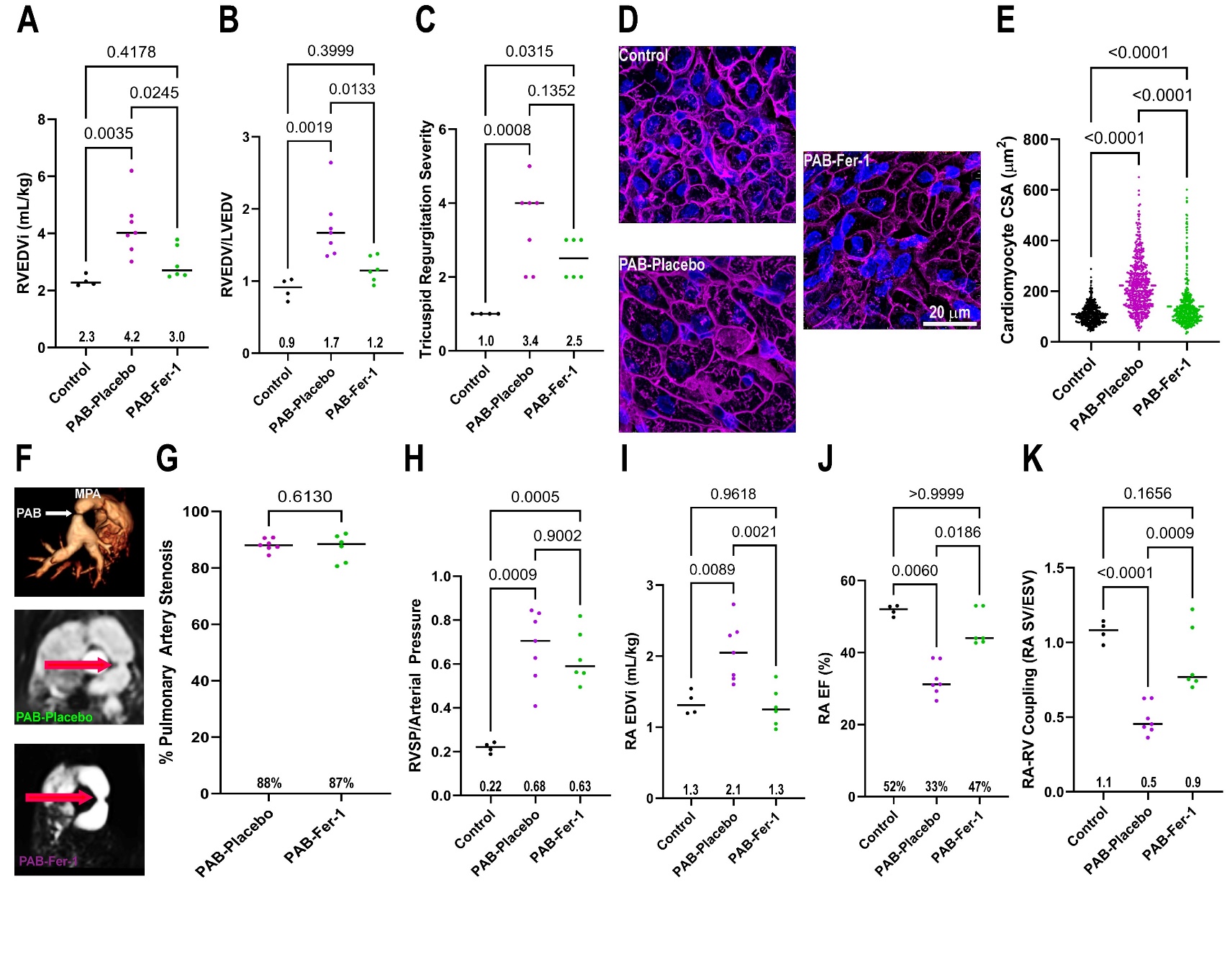

Supplemental Table 1: Results of comprehensive cardiac MRI examination

|  | Control (*n*=4) | PAB-Vehicle  (*n*=7) | PAB-Fer-1 (*n*=6) |
| --- | --- | --- | --- |
| RA EDV_i_ (mL/kg) | 1.3±0.2 | 2.1±0.4 | 1.3±0.3 |
| RA ESV_i_ (mL/kg) | 0.6±0.1 | 1.4±0.3 | 0.7±0.2 |
| RA SV_i_ (mL/kg) | 0.7±0.1 | 0.7±0.2 | 0.6±0.1 |
| RA EF (%) | 52±2 | 33±4 | 47±5 |
| RA SV/ESV | 1.1±0.1 | 0.5±0.1 | 0.9±0.2 |
| RV EDV_i_ (mL/kg) | 2.3±0.2 | 4.2±1.0 | 2.9±0.6 |
| RV ESV_i_ (mL/kg) | 1.1±0.2 | 3.0±1.1 | 1.8±0.5 |
| RV SV_i_ (mL/kg) | 1.4±0.1 | 1.2±0.2 | 1.2±0.2 |
| RV EF (%) | 59±2 | 30±7 | 41±5 |
| LV EDV_i_ (mL/kg) | 2.7±0.4 | 2.4±0.2 | 2.6±0.4 |
| LV ESV_i_ (mL/kg) | 1.1±0.3 | 1.3±0.2 | 1.2±0.2 |
| LV SV_i_ (mL/kg) | 1.5±0.2 | 1.2±0.1 | 1.3±0.2 |
| LV EF (%) | 53±3 | 48±5 | 52±3 |
| RVEDV/LVEDV | 0.9±0.2 | 1.7±0.4 | 1.2±0.2 |
| RV SV/ESV | 1.3±0.2 | 0.4±0.1 | 0.7±0.1 |
| TR Severity | 1±0 | 3.4±1.1 | 2.5±0.5 |
| Proximal PA CSA | NA | 2.3±0.3 | 2.5±0.5 |
| PA-Band CSA | NA | 0.3±0.03 | 0.3±0.1 |
| % Stenosis | NA | 88±2% | 87±5% |
| LGE Present at IVS | 0/4 | 6/7 | 3/6 |
